# High-Spatiotemporal-Resolution Nanopore Sequencing of SARS-CoV-2 and Host Cell RNAs

**DOI:** 10.1101/2021.08.20.457128

**Authors:** Dóra Tombácz, Ákos Dörmő, Gábor Gulyás, Zsolt Csabai, István Prazsák, Balázs Kakuk, Ákos Harangozó, István Jankovics, Béla Dénes, Zsolt Boldogkői

## Abstract

Recent studies have disclosed the genome, transcriptome and epigenetic compositions of severe acute respiratory syndrome coronavirus 2 (SARS-CoV-2) and the effect of viral infection on gene expression of the host cells. It has been demonstrated that, besides the major canonical transcripts, the viral genome also codes for non-canonical RNA molecules. While the structural characterizations have revealed a detailed transcriptomic architecture of the virus, the kinetic studies provided poor and often misleading results on the dynamics of both the viral and host transcripts due to the low temporal resolution of the infection event and the low virus/cell ratio (MOI=0.1) applied for the infection. In this study, we used direct cDNA and direct RNA nanopore sequencings for the generation of high-coverage, high-temporal-resolution transcriptomic datasets on SARS-CoV-2 and on primate host cells infected with a high virus titer (MOI=5). Sixteen sampling time points ranging from 1 to 96h with a varying time resolution and three biological replicates were used in the experiment for both the infected and the non-infected cells.

## Background & Summary

Severe acute respiratory syndrome coronavirus 2 (SARS-CoV-2) is a positive-sense single RNA-stranded betacoronavirus and the etiological agent of the current COVID-19 pandemic^1^. The replication and the transcription of the RNA genome are interrelated because the same enzyme, an RNA-dependent RNA polymerase (RdRP), carries out both processes^2^. First, negative-sense RNA intermediates are generated to serve as templates for the synthesis of both the genomic RNA (gRNA) and the nested set of subgenomic RNAs (sgRNAs) mapping to the 3’-third of the viral genome^3^. The gRNA and the sgRNAs have common 3’-termini since the RdRP synthesizes the positive sense RNAs from this end of the genome. Template-switching occurs during the synthesis of the negative-strand of sgRNAs, which is mediated by the transcription-regulating sequences (TRSs) in the genome <u>b</u>ody (TRS-B) and in the 5’-<u>l</u>eader sequence (TRS-L) resulting in the fusion of leader-body sequences^4,5^. Recent studies have disclosed the transcriptomic architecture of SARS-CoV-2 and the effect of viral infection on the host gene expression^6^. It has been shown that, besides canonical TRS-dependent RNA molecules, the viral genome also codes for non-canonical, TRS-dependent and TRS-independent, transcripts, although in a relative low abundance (altogether <10%). Additionally, investigations of the effect of the viral infection on the transcriptome of various cell types have identified several genes and gene networks^7^.

Nonetheless, the kinetic studies of gene expressions used only a few timepoints for monitoring the infection^8,9^, which do not provide a comprehensive picture on the temporal dynamics of viral transcriptome. Furthermore, typically a low (0.1) multiplicity of infection (MOI) was applied in the experiments, which may lead to misleading conclusions on the kinetic properties of SARS-CoV-2 transcripts, because after a couple of hours the virus can initiate a new infection cycle within the non-infected cells. Low-MOI-infection makes also difficult to assess the host cell response, especially in the case of the down-regulated genes. Infections are typically carried out using fresh, rapidly growing cells, and fresh cultures are also used as mock-infected cells. However, gene expression profiles may undergo alterations non-infected cells during the propagation therefore, we cannot decide whether the transcriptional changes in infected are due to the effect of the virus or to the time factor of culturing. This phenomenon is practically never tested in the experiments. An additional problem is the use of short-read sequencing for profiling of the host cell reaction to the viral infection^7^ because this approach has severe limitations for the detection of transcript isoforms, such as splice and length variants, and multigenic transcripts, among others^10,11,12^.

Long-read sequencing (LRS) opened new avenues for the comprehensive analysis of the transcriptomes, for which the major reason is that these techniques are able to detect full-length RNA molecules and thereby to distinguish between transcript isoforms and transcriptional overlaps. LRS-based studies have revealed a hidden transcriptional complexity in viruses^13,14^, but this approach has also been used for the analysis of the kinetic properties of viral transcriptomes^15^, and the virus-host interaction^16,17^.

In this study, we applied nanopore sequencing based on direct RNA (dRNA) and direct cDNA (dcDNA) approaches for the generation of transcriptomic datasets from SARS-CoV-2 and primate host (Vero) cells. A mixed time point sample was used for dRNA sequencing, while we used 16 time-point samples within an interval of 1 to 96 h from both infected and non-infected host cells using MOI=5 for the infection. Altogether, we generated almost 64 million long-reads, from which more than 1.8 million reads mapped to the SARS-CoV-2 and almost 48 million to the host reference genome, respectively (**Table 1**). The obtained read count resulted in a very high coverage across the viral genome (**Figure 1**). Detailed data on the read counts, quality of reads including read lengths (**Figure 2**), insertions, deletions, as well as mismatches are summarized in **Supplementary Table 1 and 2**.

**Table 1.**
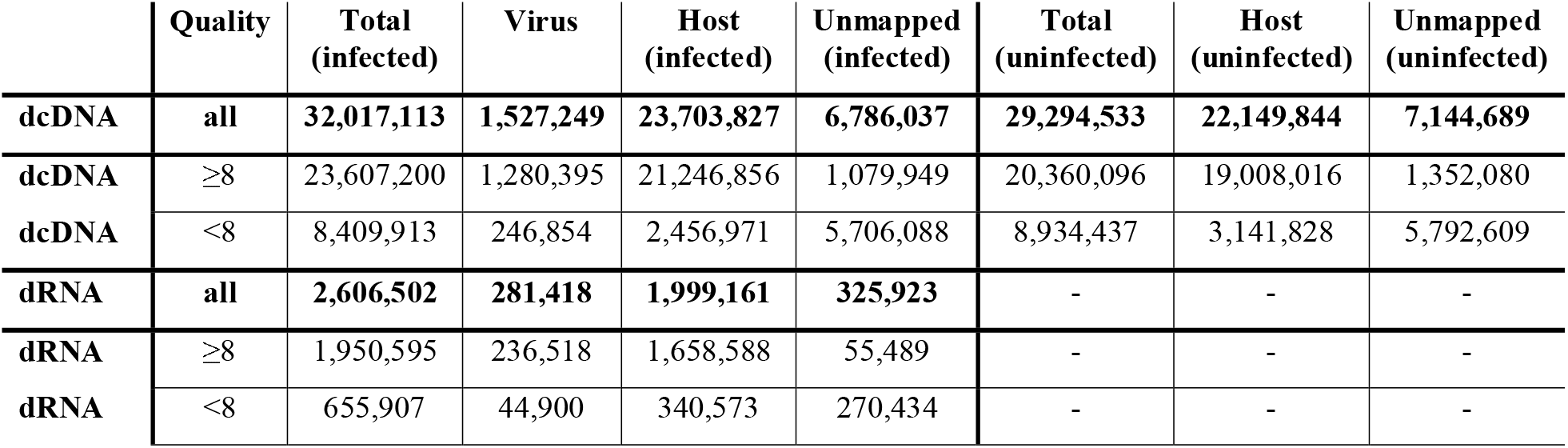
Summary data of the obtained read counts from dcDNA and dRNA sequencings. Low quality (failed) reads (Q-score <8) were filtered out from the passed reads (Q-score ≥8) by the MinKNOWs (Guppy, ONT) software.

|  | Quality | Total (infected) | Virus | Host (infected) | Unmapped (infected) | Total (uninfected) | Host (uninfected) | Unmapped (uninfected) |
| --- | --- | --- | --- | --- | --- | --- | --- | --- |
| **dcDNA** | **all** | **32,017,113** | **1,527,249** | **23,703,827** | **6,786,037** | **29,294,533** | **22,149,844** | **7,144,689** |
| **dcDNA** | ≥8 | 23,607,200 | 1,280,395 | 21,246,856 | 1,079,949 | 20,360,096 | 19,008,016 | 1,352,080 |
| **dcDNA** | <8 | 8,409,913 | 246,854 | 2,456,971 | 5,706,088 | 8,934,437 | 3,141,828 | 5,792,609 |
| **dRNA** | **all** | **2,606,502** | **281,418** | **1,999,161** | **325,923** | - | - | - |
| **dRNA** | ≥8 | 1,950,595 | 236,518 | 1,658,588 | 55,489 | - | - | - |
| **dRNA** | <8 | 655,907 | 44,900 | 340,573 | 270,434 | - | - | - |

**Figure 1.**
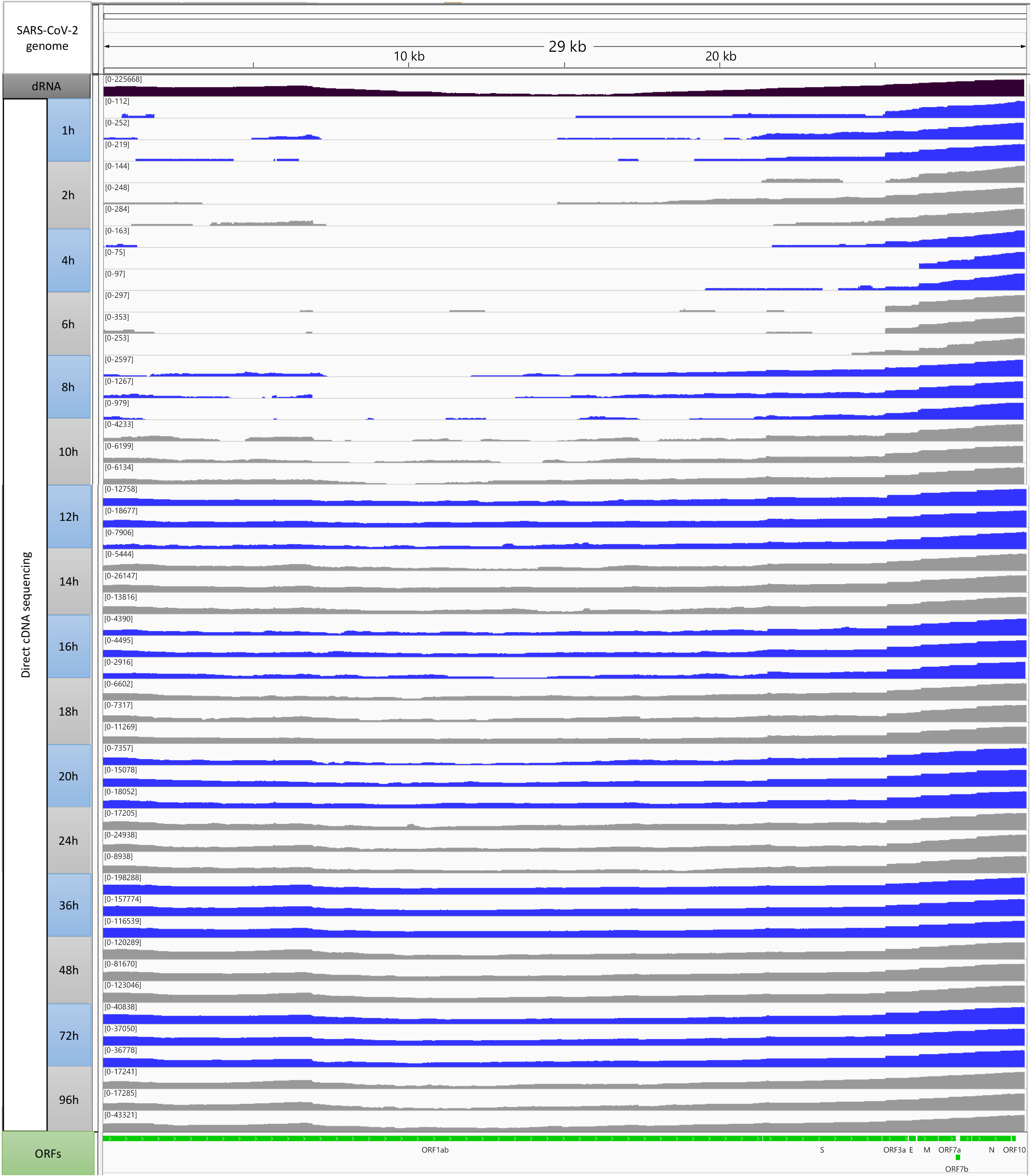
Whole genome coverage plot using high-quality (Q-score ≥8) reads which aligned to the SARS-CoV-2 genome used as references for this study.

**Figure 2.**
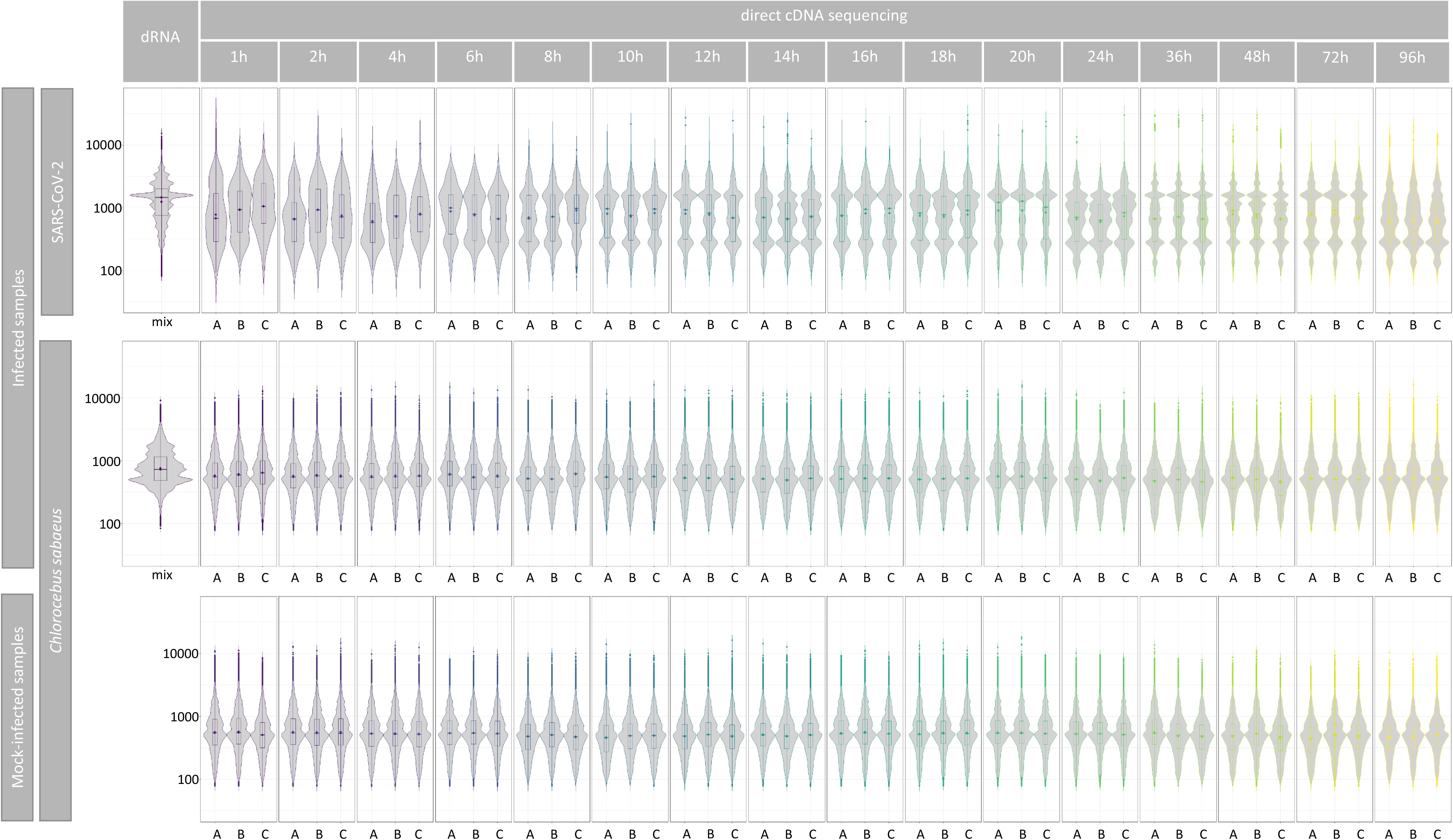
Violin plots of read length distribution of the sequencing data derived from infected and uninfected samples. **a**. Length of reads aligned to the viral **b**. and to the host genome. **c**. Read-length distribution of mock-infected samples mapped to the host genome.

Decoding the transcriptional landscape of SARS-CoV-2 virus is a fundamental step in studying its biology, genetics regulation and molecular pathogenesis. Therefore, in this data descriptor, our aim was to provide a robust, precise, reliable dataset based on LRS approaches for understanding the gene expression and genetic regulation of the causative agent of current pandemic, its effect on differential host gene expression, as well as to provide a rich resource for future functional studies.

## Methods

**Figure 3** shows the detailed workflow of the study.

**Figure 3.**
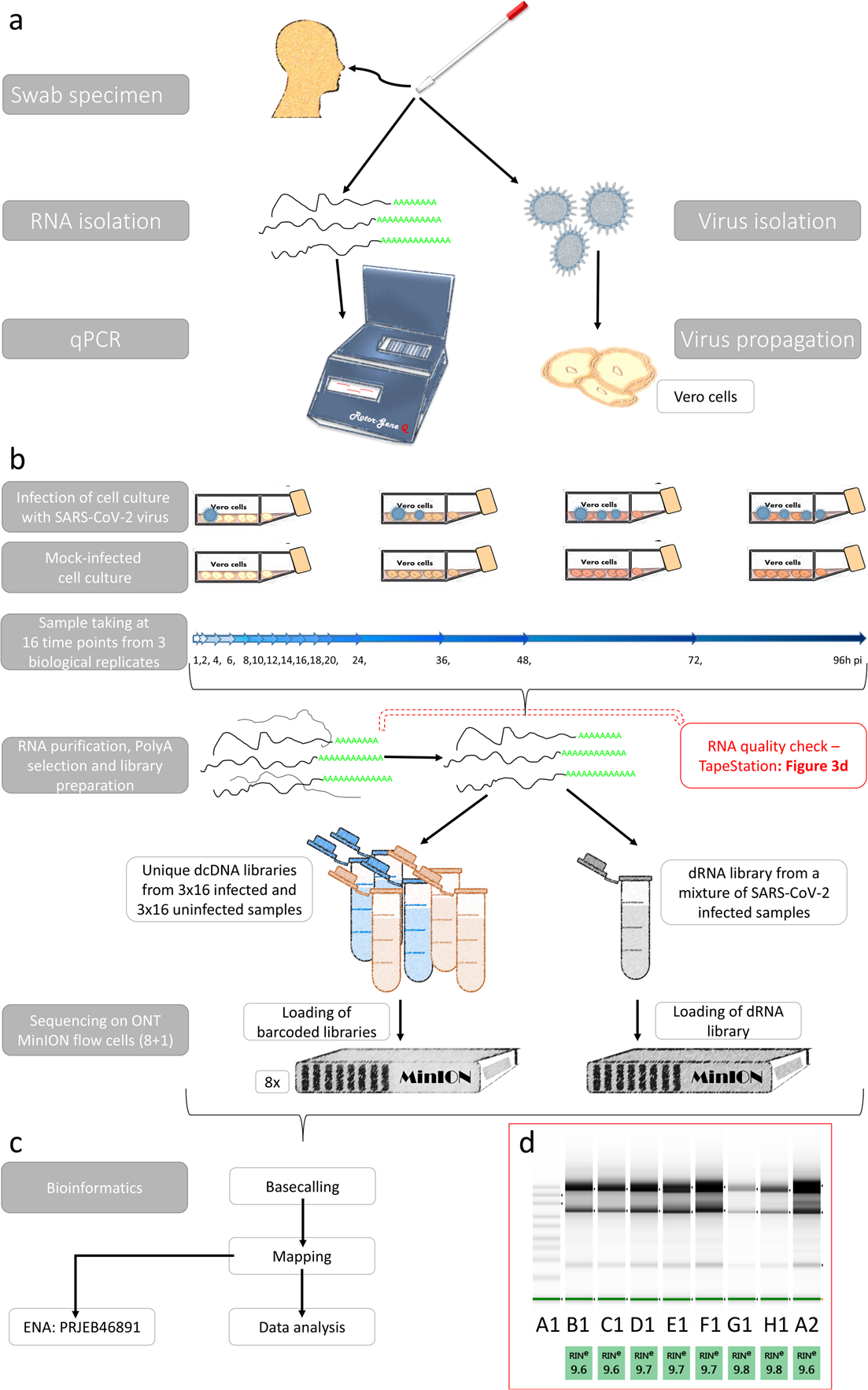
Schematic representation of the workflow applied in this project. **a**. Isolation and detection of a Hungarian isolate of SARS-CoV-2 virus. **b**. Experimental workflow of the study. **c**. Bioinformatics workflow **d**. Quality of RNA samples were detected with a TapeStation 2200 system with RNA ScreenTape. TapeStation gel image shows that intact, high quality RNAs were isolated from the samples and used for sequencing. The image shows the following samples: A1: marker; B1: 8h pi sample C; 12h pi sample A; 16h pi sample A; 18h pi sample B, 20h pi sample C; 36h pi sample A; 48h pi sample A; 96h pi sample B

### Cells

The Vero E6 (African green monkey kidney) cell line was obtained from the American Type Culture Collection (ATCC). The cells were plated at a density of 2 × 10^6^ cells per 75 cm^2^ tissue culture flasks (CELLSTAR® Greiner Bio-One GmbH, Frickenhausen, Germany) in Minimum Essential Medium Eagle culture medium (MEM) with 10% fetal bovine serum (FBS) and 2mM L-glutamine and antibiotic-antimycotic solution (all obtained from Sigma-Aldrich). Vero cells were incubated at 37°C in a humidified 5% CO_2_ atmosphere until confluency (~8 × 10^6^ cells) was reached. The monolayer was washed once with the serum-free MEM immediately before infection.

### Collection, detection and isolation of the virus

The SARS-CoV-2 virus was isolated from the human nasopharyngeal swab of the RT-PCR positive (Ct 22) 77-year-old male patient during the official COVID-19 surveillance program at the Veterinary Diagnostic Directorate of the National Food Chain Safety Office (Budapest, Hungary) with the cooperation of the Complex Medical Center (Budapest) in November 2020 at the second wave of COVID-19 pandemic in Hungary. The patient developed respiratory illness, with fever, cough, and fatigue that quickly progress to pneumonia. The patient was hospitalized, where, unfortunately, he died in a few days. In his story, he did not declare any travel abroad in the last 14⍰days. At the same time, he traveled relatively frequently within Hungary and had been in close contact with the people with COVID-19.

Detection of SARS-CoV-2 in pharyngeal wash samples was performed using RT-PCR amplification of SARS-CoV-2 N-gene fragments. Two hundred microliters (200 μL) of the pharyngeal washes were first processed for RNA extraction in the Thermo Scientific™ KingFisher™ Flex Purification System (Thermo Fisher Scientific, Waltham, MA USA), using the IndiMag® Pathogen Kit (QIAGEN® GmbH, Hilden, Germany). Subsequently, the detection of N-gene of SARS-CoV-2 was performed by using the 2019-nCoV-2 RUO kit (Integrated DNA Technologies, Inc., Coralville, Iowa, USA) and One-Step RT-PCR Kit (QIAGEN® GmbH) on a Rotor-Gene Q real-time PCR cycler (QIAGEN® GmbH). The amplification protocol consisted of a reverse transcription step at 50°C for 30 minutes, a denaturation step at 95°C for 15 minutes and subsequent 45 cycles at 95°C/56°C/72°C for 30/30/60 seconds, respectively. A positive result was defined as amplification of N-gene in a sample with each cycle threshold value (ct) less than 37.

For the virus isolation, 1 ml of viral transport media from the swab was mixed with 3 ml serum-free MEM culture medium supplemented with 2mM L-glutamine and antibiotic-antimycotic solution and were filtered using Ministar® 0.22μm filter (Sartorius AG, Göttingen, Germany). The filtrate was placed onto cells in a 25 cm^2^ tissue culture flask (Corning®, Corning Inc., New York, USA) of Vero E6 cells, then were incubated at 37°C in a humidified 5% CO2 chamber for 1 hour. After incubation, two and a half milliliters (2.5 ml) of serum-free MEM culture medium with 2mM L-glutamine and antibiotic-antimycotic solution was added to the tissue culture flask. The inoculated culture was grown in a humidified 37°C incubator with 5% CO2. Cells were observed daily for cytopathic effect (CPE). On day 4, an 80% CPE was evident, and the cells with supernatant were harvested. This provided the first passage virus. The virus was passaged twice at low MOI in Vero E6 cells to obtain a working stock used in the experiments. The viral titer was determined by plaque assay on Vero E6 cells. Virus stock was stored at −80C until use.

### Propagation of the virus

The virus was passaged twice in Vero cells to obtain a working stock used in all experiments. Viral titer was determined by plaque assay on Vero cells. The virus was diluted into a serum-free MEM. Cells were infected with 5 ml of the SARS-CoV-2 virus with 5 plaque-forming unit (pfu)/cell [multiplicity of infection (MOI=5)], then were incubated at 4°C for 1 h. Non-infected control cultures (mock) were prepared using pure non-supplemented MEM as inoculums. Next, the virus inoculum was removed from the flasks. The monolayer was washed once with the serum-free MEM. Ten milliliters of MEM culture medium supplemented with 3% FBS, 2mM L-glutamine, and antibiotic-antimycotic solution was added to the tissue culture flasks. The cells were incubated at 37°C for 1, 2, 4, 6, 8, 10, 12, 14, 16, 18, 20, 24, 36, 48, 72 and 96 hours in a humidified 5% CO_2_ atmosphere. Each time experiment was done in triplicate with a mock-infected control. Mock-infected cells were harvested at the same time points as the infected cells. Following incubation, the medium was removed, and the monolayer was washed once with phosphate-buffered saline (PBS). The tissue culture plates were stored at −80°C until use. Next, the infected cells were treated by lysis buffer, then creped and placed into an Eppendorf Tubes^®^ (Thermo Fisher Scientific Inc.).

### RNA purification

Total RNA was extracted from the mock-infected and from the SARS-CoV-2-infected cells at various stages of infection from 1 to 96 hours using the Macherey-Nagel’s NucleoSpin RNA Kit according to the manufacturer’s protocol. In brief, cells were collected by low-speed centrifugation, then 350μl lysis buffer (RA1 from the Kit), 3.5μl β-Mercapthoethanol (Sigma Aldrich) were added followed by vortexing the samples. Mixtures were loaded onto a NucleoSpin Filter and centrifuged for 1min at 11,000 x g. The filters were discarded and 350μl 70% EtOH was added to the lysate. This was loaded to the NucleoSpin RNA Column and centrifuged at 11,000 x g for 30sec. Membrane was desalted with the addition of 350μl Membrane Desalting Buffer (from the NucleoSpin Kit), then dried with a short centrifugation (11,000 x g). Residual DNA was enzymatically removed [with the usage of 95μl mixture of rDNase:rDNase reaction buffer (1:9 ratio, NucleoSpin Kit) and incubation at room temperature (RT) for 15min]. The rDNase was inactivated with the first washing step, by adding 200μl RAW2 Buffer (NucleoSpin Kit) directly onto the NucleoSpin Filter. After a quick centrifugation (30min, 11,000 x g), the filter was placed in a new tube. Six-hundred μl RAW3 Buffer (NucleoSpin Kit) was added, then spun down as before. This washing step was repeated using 250μl RAW3. Finally, the total RNA bound to the Filter was eluted in 60μl nuclease-free water (NucleoSpin Kit). Samples were quantified by Qubit 4.0 using the Qubit RNA BR Assay Kit (Invitrogen, **Supplementary Table 3a**) and then stored at −80°C until use.

### Poly(A) selection

Lexogen’s Poly(A) RNA Selection Kit V1.5 was used to isolate polyadenylated RNAs from the total RNA samples. The protocol applies oligo(dT) beads, which capture RNAs with poly(A) stretches (most mRNAs), but RNAs without polyadenylated 3’ ends (e.g. 28S and 18S rRNAs and tRNAs) do not hybridize to the beads and therefore, they will be removed during the washing steps. The detailed protocol is as follows: the magnetic beads (part of the Lexogen Kit) was resuspended and 4μl for each RNA samples was measured. Beads were placed in a magnet, they were collected and the supernatant was discarded. Samples were resuspended in 75μl Bead Wash Buffer (Lexogen Kit) and then were placed on the magnet. Supernatant was discarded and this washing step was repeated once. Beads were resuspended in 20μl RNA Hybridization Buffer (part of the Lexogen Kit). Ten ng from the total RNA samples were diluted to 20μl UltraPure™ DNase/RNase-Free Distilled Water (Invitrogen) and then denatured at 60°C for 1min followed by holding them at 25°C. Twenty μl denatured RNA was mixed with 20μl (previously washed and resuspended) beads. The mixtures were incubated at 25°C in a shaker incubator with 1250 rpm agitation. After 20min incubation, sample-containing tubes were placed in a magnetic rack. Supernatant was discarded then the tubes were removed from the magnet. Samples were resuspended in 100μl Bead Wash Buffer (Lexogen Kit) then they were incubated for 5min at 25°C with 1250 rpm agitation. Supernatant was discarded and the washing step was repeated. After the complete removal of the supernatant, beads were resuspended in 12μl UltraPure™ DNase/RNase-Free Distilled Water. Samples were incubated at 70°C for 1min, then the tubes were placed on a magnetic rack. Supernatant, containing the poly(A)+ RNA fraction, was placed to new DNA LoBind (Eppendorf) tubes, the RNA concentration was measured using Qubit RNA HS Assay Kit (Invitrogen, **Supplementary Table 3b**), then samples were stored at −80°C.

### ONT – direct cDNA sequencing

For the analysis of the dynamic properties of SARS-CoV-2 RNAs and the effect of viral infection on the host cell transcriptome profile, RNA samples from different time points (1, 2, 4, 6, 8, 10, 12, 14, 16, 18, 20, 24, 36, 48, 72 and 96h p.i., **Supplementary Table 3c**) were used individually for the generation of direct cDNA libraries for nanopore sequencing. The non-amplified cDNA libraries were prepared from sixteen time-points from the mock, and from the coronavirus-infected samples, in three biological replicates using the Direct cDNA Sequencing Kit (SQK-DCS109, ONT) and the appropriate ONT protocol. In short, first-strand cDNAs were generated from the polyA(+) RNAs using the Maxima H Minus Reverse Transcriptase (Thermo Fisher Scientific) with SSP and VN primers (supplied in the kit). The RNase Cocktail Enzyme Mix (Thermo Fisher Scientific) was used to eliminate the potential RNA contamination. Synthesis of the second cDNA strands were carried out with LongAmp Taq Master Mix (New England Biolabs). The double-stranded cDNAs were repaired (NEBNext End repair /dA-tailing Module, New England Biolabs) and adapter ligated (NEB Blunt /TA Ligase Master Mix New England Biolabs). Individual barcode sequences were added to each samples for multiplex sequencing for which the Native Barcoding (12) Kit (ONT) was used as recommended by the manufacturer. The cDNAs and the libraries were washed using AMPure XP beads (Agencourt, Beckman Coulter) after every enzymatic reaction step. The barcode labeled samples were loaded onto MinION R9.4 SpotON Flow Cells (ONT, **Table 2**).

**Table 2.**
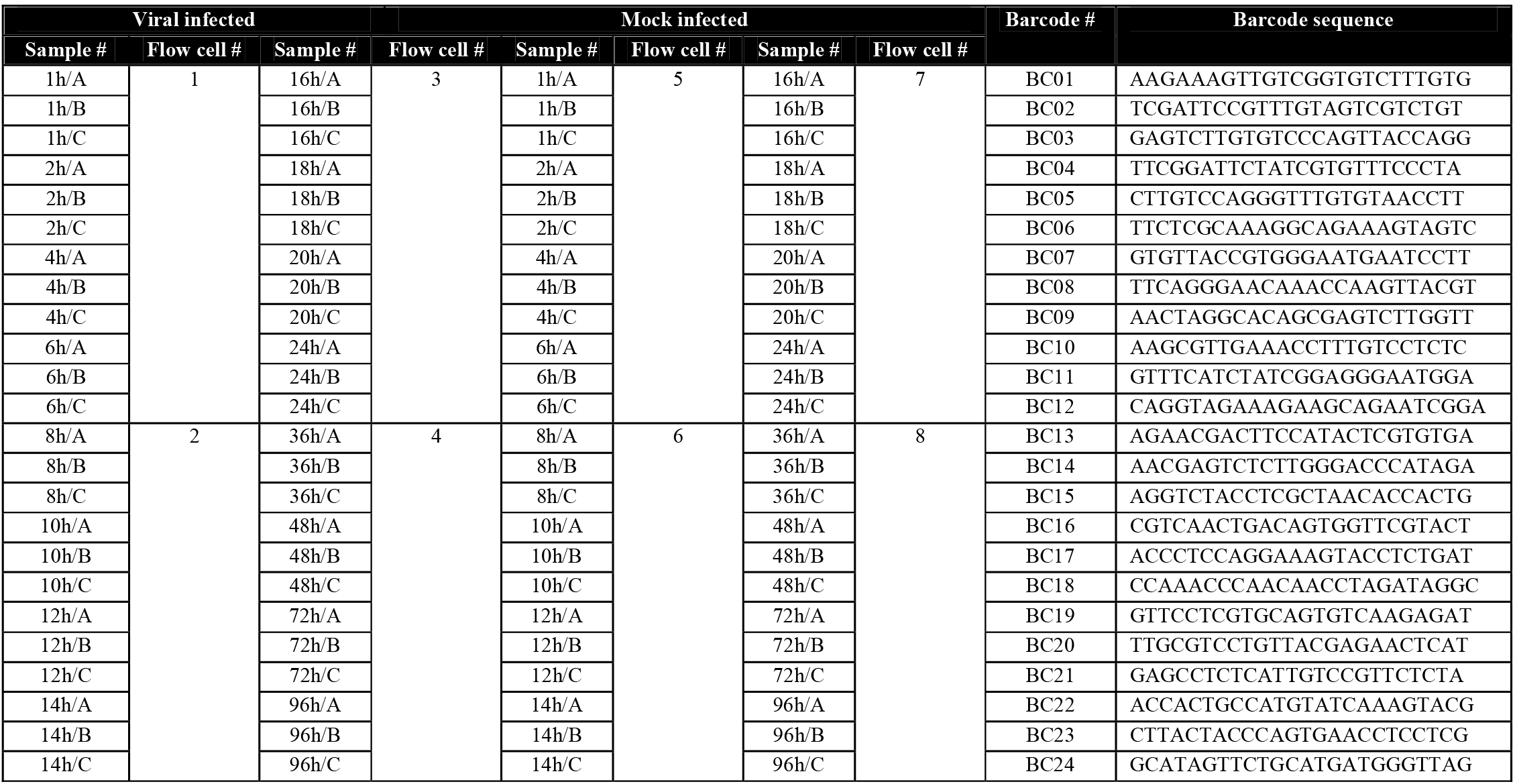
List of the sequences of barcodes used for multiplex sequencing. This table also contains the information about the barcoded samples loaded on the same flow cell. A, B, and C represent the biological replicates.

### ONT – direct RNA sequencing

ONT’s Direct RNA sequencing (SQK-RNA002; Version: DRS_9080_v2_revO_14Aug2019, Last update: 10/06/2021) was used to sequence the native RNA strands from a mixture of polyA(+) RNA fractions (**Supplementary Table 3d**). Five-hundred ng RNA in 9μl nuclease-free water was mixed with 3μl NEBNext Quick Ligation Reaction Buffer (New England BioLabs), 0.5μl RNA CS (ONT Kit), 1μl RT Adapter (110nM; ONT Kit) and 1.5μl T4 DNA Ligase (2M U/ml New England BioLabs). The ligation reaction was carried out for 10min at RT. The synthesis of the first strand cDNA was conducted using SuperScript III Reverse Transcriptase (Life Technologies), as described in the Direct RNA sequencing (DRS) protocol (ONT). In short, a 50 min incubation at 50°C was followed by the inactivation of the enzyme at 70°C for 10 min. Sequencing adapters from the DRS kit were ligated to the cDNA with the T4 DNA ligase enzyme and NEBNext Quick Ligation Reaction Buffer (New England BioLabs). Ligation was carried out at RT for 10 min. The sample was washed using AMPure XP beads (Agencourt, Beckman Coulter) after every enzymatic reaction. Libraries were sequenced on an R9.4 SpotON Flow Cell.

### Pre-processing and data analysis

The MinION raw data was basecalled using ONT Guppy basecalling software version 5.0.11. using -- qscore_filtering. VirStrain^18^ tool was used to identify the closest SARS-CoV-2 strains to our isolate (**Supplementary Table 4**). Then, reads with a Q-score greater or equal to 8 were mapped to the SARS-CoV-2 viral genome (NCBI nucleotide accession: MT560672.1) and to the host (Chlorocebus_sabeus 1.1) genome [GenBank assembly accession: GCA_000409795.2 (latest)] with the Minimap2 aligner^19^. The mock-infected samples were also mapped to the host genome. The following programs were also used for analysis: SamTools^20^ [for generation binary alignment (bam) and indexed (bai) files, as well as to categorize the data into viral-mapped, host-mapped and unmapped], the bamCoverage tool from deepTools^21^ (to generate coverage tracks), GATK^22^ Picard’s DownsampleSam tool which applies a downsampling algorithm to retain only a deterministically random subset of the reads. R package^23^ (R script) was used for generate violin plots.

Our in-house scripts were used to generate the descriptive statistics of reads: https://github.com/moldovannorbert/seqtools.

## Data Records

All data generated in this study including the unmapped reads as well as reads which do not match our strict criteria (Q-score below 8) can be found in European Nucleotide Archive under the accession number: **PRJEB46891**. Data records and supporting information (e.g. sample IDs, experimental steps performed on each, etc.) are summarized in **Supplementary Table 5**.

## Technical Validation

### RNA

The Qubit RNA BR Assay Kit (Invitrogen, Carlsbad, CA, United States) was used to check the amount of total RNA. Qubit RNA HS Assay Kit (Invitrogen, Carlsbad, CA, United States) was used to measure the quantity of the poly(A)+ RNA fractions. The final concentrations of the RNA samples were determined by Qubit^®^ 4.

### cDNA

The concentrations of the cDNA samples and sequencing ready libraries were measured using the Qubit dsDNA HS Assay Kit (Invitrogen, Carlsbad, CA, United States). The quality of RNA was assessed using the Agilent 2200 TapeStation System. RIN scores ≥ 9.6 were used for sequencing (**Figure 3d**).

The cDNAs and the sequencing-ready cDNA libraries were washed using AMPure XP beads (Agencourt, Beckman Coulter) after every enzymatic reaction. The samples for dRNA sequencing were treated with RNAClean XP beads.

Three biological replicates were used for each of the 16 time points. To monitor the effect of SARS-CoV-2 infection on the gene expression of the host cells, mock-infected cells were harvested at the same time-points, as the virally-infected cells.

## Usage Notes

The datasets provided in this report are useful to investigate the time-course expression of SARS-CoV-2 over a 96 h period of infection which provides a deeper understanding of the molecular biology of the virus (e.g. transcriptional analysis of subgenomic region, analysis of the dynamics of viral replication, examination of the potential interactions between transcription and replication, as well as to study the potential transcript isoforms of the virus). Our data eliminate the limitations of other SARS-CoV-2 transcriptomic experiments. First, we used a high plaque-forming unit per cell (MOI=5 pfu/cell) for the infection (other studies typically apply 0.1 pfu), therefore the large majority of cells in the culture became infected, and hence the possibility of a second round of infection is excluded. Additionally, due to the high temporal resolution, our data is also useful to precisely measure the alteration of the gene expression of both the virus and the host cell. Third, we provide mock-infected cells, which were harvested in the same time-points, as the virally-infected cells, which allows the identification of gene-network alterations due to the aging of the cell culture and to analyze the temporal changes of gene expression patterns during the cultivation. Virus-host interactions can also be examined. Furthermore, due to the very long reads and high coverage across the viral genome, assembly of this Hungarian isolate and the analysis of potential genome editing events can be achieved from the data. Moreover, the applied direct RNA and direct cDNA sequencing approaches provides independent methods for the validation of novel transcripts. Due to the high coverage, this dataset can also be used for the reconstruction of the viral genome. Finally, this dataset can also be used from various bioinformatics aspects: e.g. the data can be further analyzed or other bioinformatic programs [NanoPack^24^, SQANTI3^25^ (https://github.com/ConesaLab/SQANTI3), lra^26^, LoRTIA^27^ (https://github.com/zsolt-balazs/LoRTIA) or any other programs for LRS data analysis listed in LONG-READ-TOOLS^28,29^] can be tested on these data. Potential template switching artefacts can be tested using the transcript annotator developed by our group^27^.

The uploaded binary alignment (BAM) files contain reads already mapped to the SARS-CoV-2 reference genome (MT560672.1), as well as to the host genome (GCA_000409795.2) using Minimap2.

The uploaded reads contain terminal poly(A) sequences as well as the 5⍰ and 3⍰ adapters, which can be used to determine the orientations of the reads.

## Code Availability

1. Custom (in-house) scripts: https://github.com/moldovannorbert/seqtools
2. deepTools: https://deeptools.readthedocs.io/en/develop/content/tools/bamCoverage.html?highlight=bamcoverage
3. GATK: https://gatk.broadinstitute.org/hc/en-us/articles/4404604771227-DownsampleSam-Picard-
4. Guppy: https://github.com/nanoporetech/pyguppyclient
5. Minimap2: https://github.com/lh3/minimap2
6. R package: https://www.r-project.org/
7. SamTools: https://github.com/SAMPTools/
8. VirStrain: https://github.com/liaoherui/VirStrain

## Acknowledgements

This project was supported by the National Research, Development and Innovation Office (OTKA) K 128247 to ZB, National Research, Development and Innovation Office (OTKA) FK 128252 to DT. This research was also supported by the project “Professional, methodological development of health care system: subproject E – Complex epidemiological and clinical response to Covid-19 pandemic “(EFOP-1.8.0-VEKOP-17-2017-00001) to BD. The APC fee was covered by the University of Szeged Open Access Fund: 5401.

## Author contributions

D.T. analyzed the data, took part in Nanopore sequencing, drafted the manuscript and coordinated the project. Á.D. performed Nanopore sequencing and RNA purification. G.G conducted bioinformatics analysis. Z.C. took part in RNA isolation and sequencing. I.P. participated in RNA isolation and analysis. B.K. carried out bioinformatics. Á.H. participated in sequencing and data analysis. I.J. isolated and propagated the virus. B.D. propagated the virus and the host cells and took part in drafting the manuscript. Z.B. conceived and designed the experiments, supervised the project and wrote the manuscript. All authors read and approved the final paper.

